# Quaternary structures of *Streptococcus pneumoniae* Nucleoside Diphosphate Kinase: from hexamers to supramolecular assemblies

**DOI:** 10.1101/2025.10.09.681369

**Authors:** Julie Kerboeuf, Paul Nouri, Frédéric Galisson, Laetitia Daury, Marie-France Giraud, Olivier Lambert, Cédric Orelle, Lionel Ballut, Elise Kaplan, Jean-Michel Jault, Cécile Gonzalez

**Author notes:** These two authors participated equally to the work (Julie Kerboeuf, Paul Nouri).

## Abstract

The Nucleoside Diphosphate Kinase (NDK) is a key enzyme that controls the balance of nucleotide pools in all living organisms. Beyond this fundamental role, NDKs exert pleiotropic effects in many cellular processes, including cell development, signal transduction, differentiation, tumor metastasis, and gene expression. The quaternary structure of NDK is typically hexameric—organized as a trimer of dimers in eukaryotic cells and in many prokaryotes—though in some species it can also be found as a tetramer (a dimer of dimers). Here, we report the crystal structure of *Streptococcus pneumoniae* NDK (SpNDK) in its apo state (1.2 Å) and in an ADP-vanadate-bound state (3.4 Å). In both structures, SpNDK adopts a hexameric assembly and the fold of each monomer is highly conserved compared to NDKs from other organisms. A notable feature is the extended *Kpn*-loop, which plays a key role in stabilizing the hexamer. Interestingly, the thermal stability of SpNDK was remarkably high (~75 °C), a surprising feature given the mesophilic lifestyle of *S. pneumoniae*. Unexpectedly, size-exclusion chromatography revealed that SpNDK exists as an equilibrium mixture of hexamers, dodecamers, and higher-order supramolecular assemblies, and this was further confirmed by mass photometry. Using cryo-electron microscopy, we solved the 3D structure of the hexameric state at 2.47 Å resolution, and resolved dodecameric assemblies of the protein. Since the oligomeric state of the NDK influences its cellular function, our findings suggest that supramolecular assemblies could modulate the activity of SpNDK *in vivo*, aligning with the broad and multifaceted roles of this enzyme family.

## Introduction

*Streptococcus pneumoniae* is a Gram-positive bacterium responsible for a broad spectrum of infections, including pneumonia, meningitis, and sepsis^1^. While vaccines targeting some *S. pneumoniae* serotypes are available^2^, the rising prevalence of antibiotic-resistant strains poses a significant public health challenge, and underscores the urgent need for novel therapeutic strategies^3^. At the molecular level, targeting specific proteins—particularly enzymes essential for bacterial growth—offers promising avenues to combat *S. pneumoniae* infections. Moreover, proteins that contribute to bacterial virulence represent valuable therapeutic targets for curbing pathogen dissemination during infection. Among these, enzymes involved in nucleotide metabolism, such as nucleoside diphosphate kinase (NDK)^4^, are of particular interest. NDKs (EC12.7.4.6) are ubiquitous enzymes that catalyze the transfer of the terminal phosphate group from nucleoside triphosphates (NTPs) to nucleoside diphosphates (NDPs). Beyond their pivotal role in maintaining cellular nucleotide homeostasis, NDKs participate in numerous biological processes including DNA replication, signal transduction, and cellular proliferation^5,6^. In bacteria, NDKs have also been implicated in stress response, biofilm formation, and the regulation of virulence-associated pathways and host cell defense mechanisms^7–10^.

NDKs are evolutionarily conserved enzymes present across all domains of life and share a common ferredoxin-like fold at the monomeric level^11^. However, their biological assemblies differ depending on the source of the organism. In eukaryotes, including mammals and plants, NDKs typically assemble as hexamers and multiple isoforms exist, up to ten, with location spanning the cytoplasm, mitochondria, chloroplast and nucleus^11,12^. In bacteria, NDKs occur in three main oligomeric forms : hexamers are typical of Gram-positive bacteria such as *Staphylococcus aureus*^13^ and *Mycobacterium tuberculosis*^14^; tetramers can be found in some Gram-negative species like *E. coli* ^15^; and dimers are observed in others, such as *Halomonas sp*.^16^. Regardless of origin, the building block for oligomer formation is always a dimer, with tetramers and hexamers arising from the association of two or three dimers, respectively^11^.

Here, we present the first X⍰ray crystal structure of *Streptococcus pneumoniae* NDK (SpNDK) at 1.21Å resolution. The asymmetric unit of this crystal comprises a single monomer; nevertheless, crystallographic symmetry operations reconstitute the canonical hexamer, consistent with the physiological oligomeric state of the enzyme. Additionally, we determined the structure of SpNDK in complex with ADP and vanadate at 3.4 Å resolution, revealing that the nucleotide binds in a manner analogous to previously solved NDK structures. Importantly, size-exclusion chromatography and mass photometry uncovered higher-order assemblies in solution, most prominently 12-mers and 18-mers, which likely correspond to dimers and trimers of hexamers, respectively. The existence of dodecameric species was further validated by cryo⍰electron microscopy (cryo-EM) analyses. These supramolecular assemblies have not previously been described for any bacterial enzyme before, and given the diverse cellular functions of NDKs, their presence suggests new, unrecognized regulatory or scaffolding roles.

## RESULTS

### Atomic spNDK structures

We used X-ray crystallography to determine the atomic structures of SpNDK in both its unliganded (apo) form and in complex with ADP and vanadate. Data collection and refinement statistics are presented in Table S1. The apo structure, solved at the very high resolution of 1.20 Å, contains a single monomer in the asymmetric unit (Fig. 1**A**). The overall structure of SpNDK monomer adopts a ferredoxin-like fold, characterized by an anti-parallel β-sheet core (β2-β3-β1-β4) surrounded by eight α-helices (Fig. 1**A**). As typically observed in all NDKs, the catalytic histidine, H121, is located on the β4 strand. Application of crystallographic symmetry reconstructs the NDK homohexamer, arranged as a trimer of dimers (Fig. 1**B**). The contact surface area between two protomers forming a dimer is approximately 600 Å^2^, as calculated with PISA^17^. This interface primarily involves residues Val18, Gly19, Glu26, Leu35 and Phe37 from both protomers. Within a dimer, the β-strands of the two monomers align continuously, forming a β-sheet composed of eight strands in total (Fig. 1**B**). Interestingly, SpNDK has the shortest *C*-terminal segment among NDKs whose structures have been solved previously (Fig. 2). Given the notorious involvement of this region in the hexameric assembly of many NDKs^11^, suggests that additional structural elements must contribute to stabilize the hexamer formation in SpNDK. Notably, the *Kpn*-loop is extended in SpNDK (Fig. 2) and is structurally quite diverse as compared to other NDK proteins (Fig. 1**A**). In SpNDK, this loop strongly contributes to the interaction with the two other dimers forming the hexamer (Fig. S1). In particular, Arg28 from the A-subunit is a major actor providing four hydrogen bonds with residues Pro94, Gly104 and Ala107 from the neighbor E-subunit (Fig. 3; see the subunit arrangement in Fig. S1). The NDK from *M. tuberculosis* (MtNDK) is another example with a short *C*-terminus extension (Fig. 2), which also form hexamers^14^. In this case, two residues (R80 and D93) are forming a salt bridge to stabilize the physiological assembly (Fig. S1). An extended *Kpn*-loop is also present in two NDKs from *B. burgdorferi* and *P. aerophilum* (Fig. S1), both of which assemble as hexamers. In both cases, this loop contributes to the interactions within the hexamer although different residues scattered throughout the loop participate in subunit contacts. In *B. burgdorferi* NDK, however, the longer C-terminus tail also play a role in hexamer stability.

**Figure 1.**
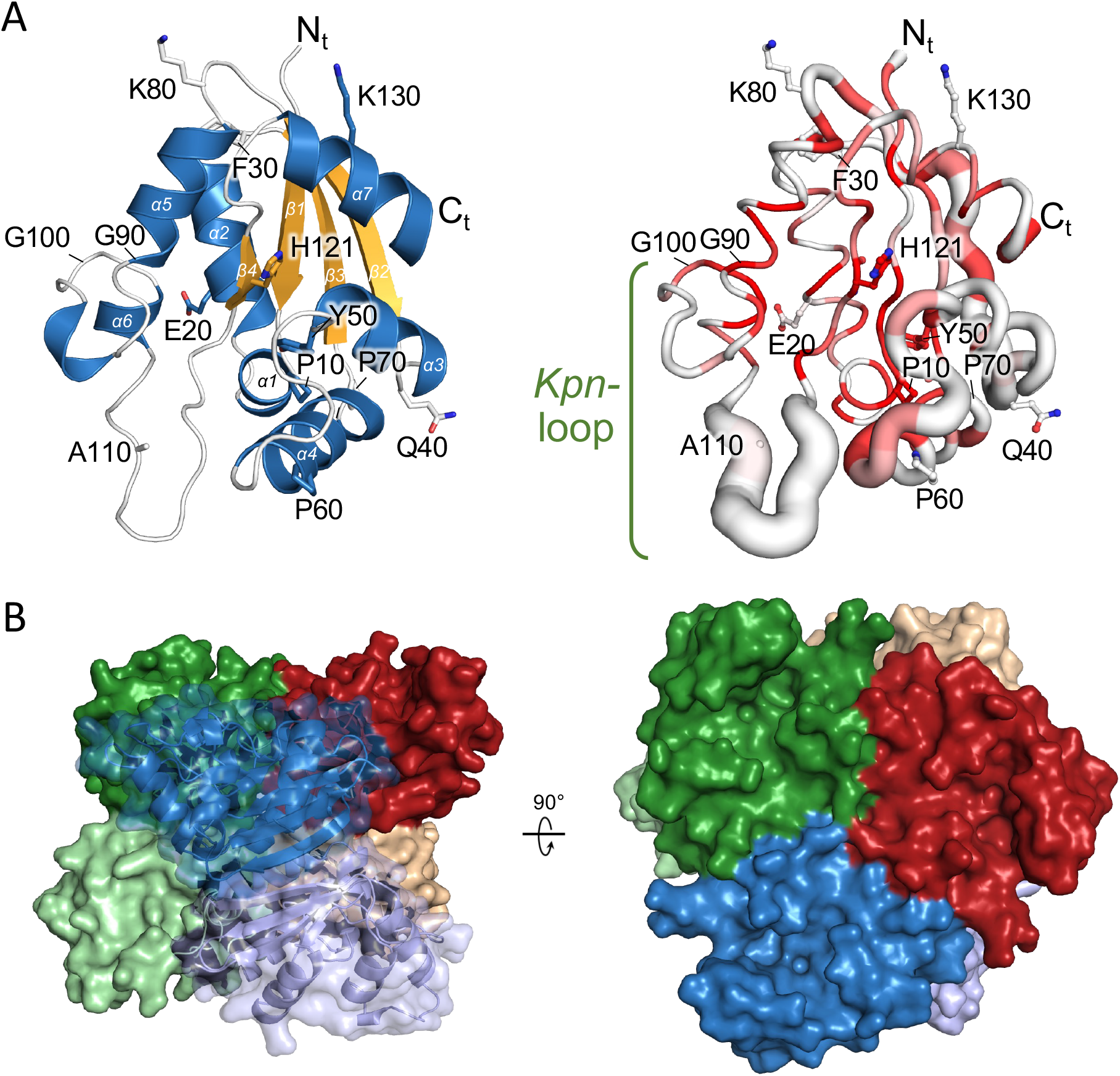
Crystal structure of the *S. pneumoniae* NDK. **A**, cartoon (left) and sausage (right) representation of the apo form of SpNDK. In the sausage representation, the protein is coloured according to sequence conservation from white, for non-conserved residues, to red for strictly conserved residues. Residues numbered at intervals of ten, except the H121 from the active site, are shown as sticks. The tube diameter is proportional to the C_α_ position difference between SpNDK and known structures of homologous NDKs using the PDBAA70 database. The right panel was drawn using ENDscript 2.0^54^ and visualized with PyMol. **B**, hexameric assembly of the apo SpNDK reconstructed from crystal symmetry. In the left panel, the dimer (blue shades) is displayed in cartoon representation with a transparent surface overlay.

**Figure 2.**
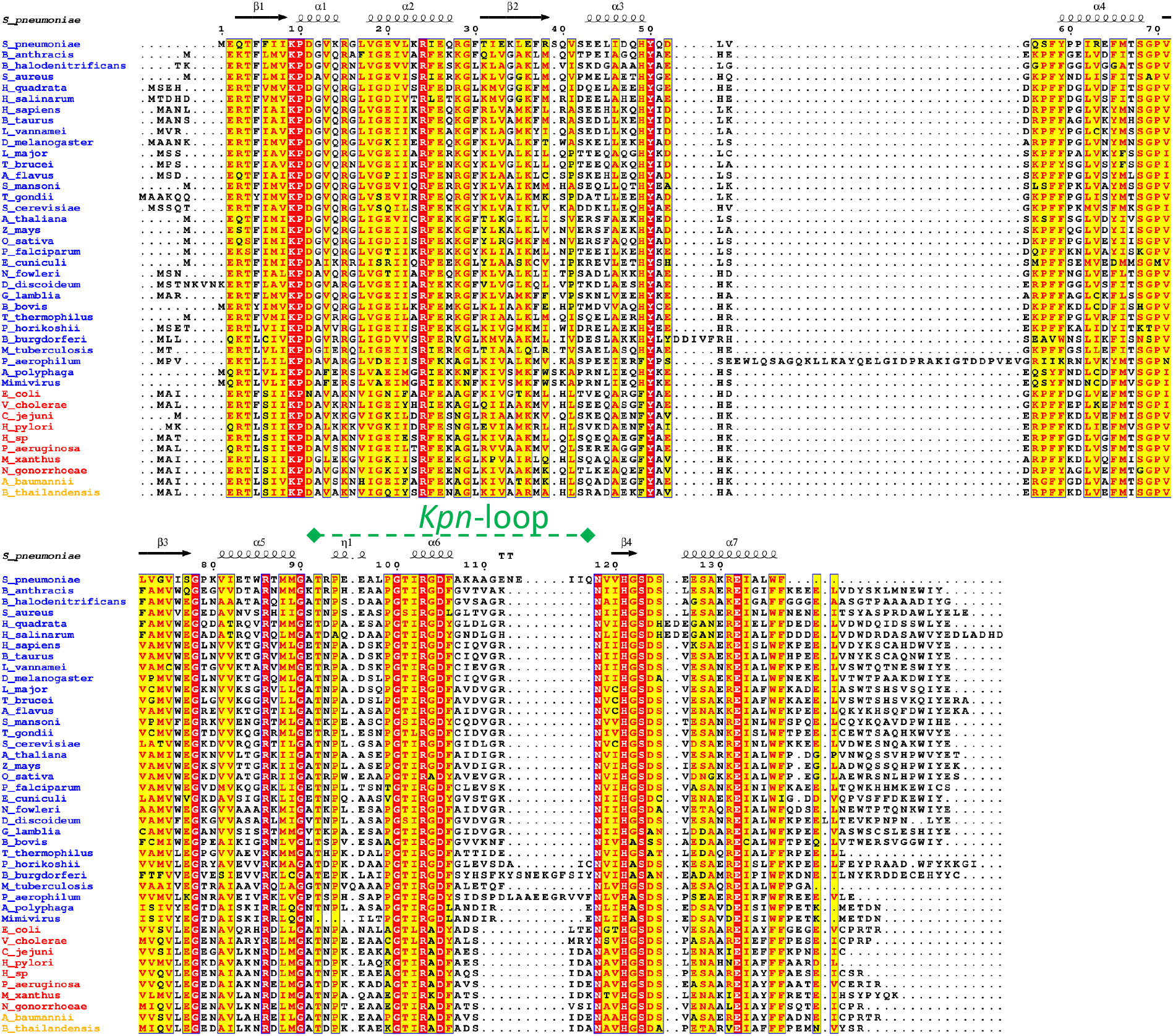
Sequence alignment of NDKs with known 3D structures. NDK sequences are from *Streptococcus pneumoniae* (PDB 9RVW), *Litopenaeus vannamei* (4UOF), *Thermus thermophilus* (1WKJ), *Schistosoma mansoni* (5IOL), *Drosophila melanogaster* (1NDL), *Aspergillus flavus* (6JOH), *Homo sapiens* (1NUE), *Arabidopsis thaliana* (1U8W), *Toxoplasma gondii* (5BXI), *Bos taurus* (1BHN), *Mycobacterium tuberculosis* (4ANC), *Oryza sativa* (1PKU), *Leishmania major* (3NGR), *Trypanosoma brucei* (4FKX), *Zea mays* (1VYA), *Giardia lamblia* (3R9L), *Saccharomyces cerevisiae* (3B54), *Naegleria fowleri* (5U2I), *Bacillus anthracis* (2VU5), *Staphylococcus aureus* (3Q83), *Acanthamoeba polyphaga (*3EM1), *Vibrio cholerae* (5X00), *Dictyostelium discoideum* (1B99), *Plasmodium falciparum* (1XIQ), *Pyrococcus horikoshii* (2CWK), *Babesia bovis* (3JS9), *Campylobacter jejuni* (3PJ9), *Haloarcula quadrata* (2ZUA), *Halobacterium salinarum* (2AZ3), *Mimivirus* (2B8Q), *Halomonas sp*. (3VGU), *Pseudomonas aeruginosa* (6AES), *Escherichia coli* (2HUR), *Acinetobacter baumannii* (4WBF), *Virgibacillus halodenitrificans* (1NB2), *Borrelia burgdorferi* (4DI6), *Myxococcus xanthus* (1NHK), *Neisseria gonorrhoeae* (5V6D), *Burkholderia thailandensis* (4DUT), *Helicobacter pylori* (6AY1), *Encephalitozoon cuniculi* (3MPD), *Pyrobaculum aerophilum* (1XQI). Species names in blue indicate NDK crystal structures that form hexamers, while those in red correspond to tetrameric assemblies. Names in orange indicate dimers, with the physiological assembly presumed to be hexameric. The figure was drawn with ESPript 3.0 with strictly conserved residues shown on a red background, and highly similar residues (present in at least 70% of the sequences) displayed on a yellow background^54^. The secondary structure of SpNDK is shown on top of the alignment.

**Figure 3.**
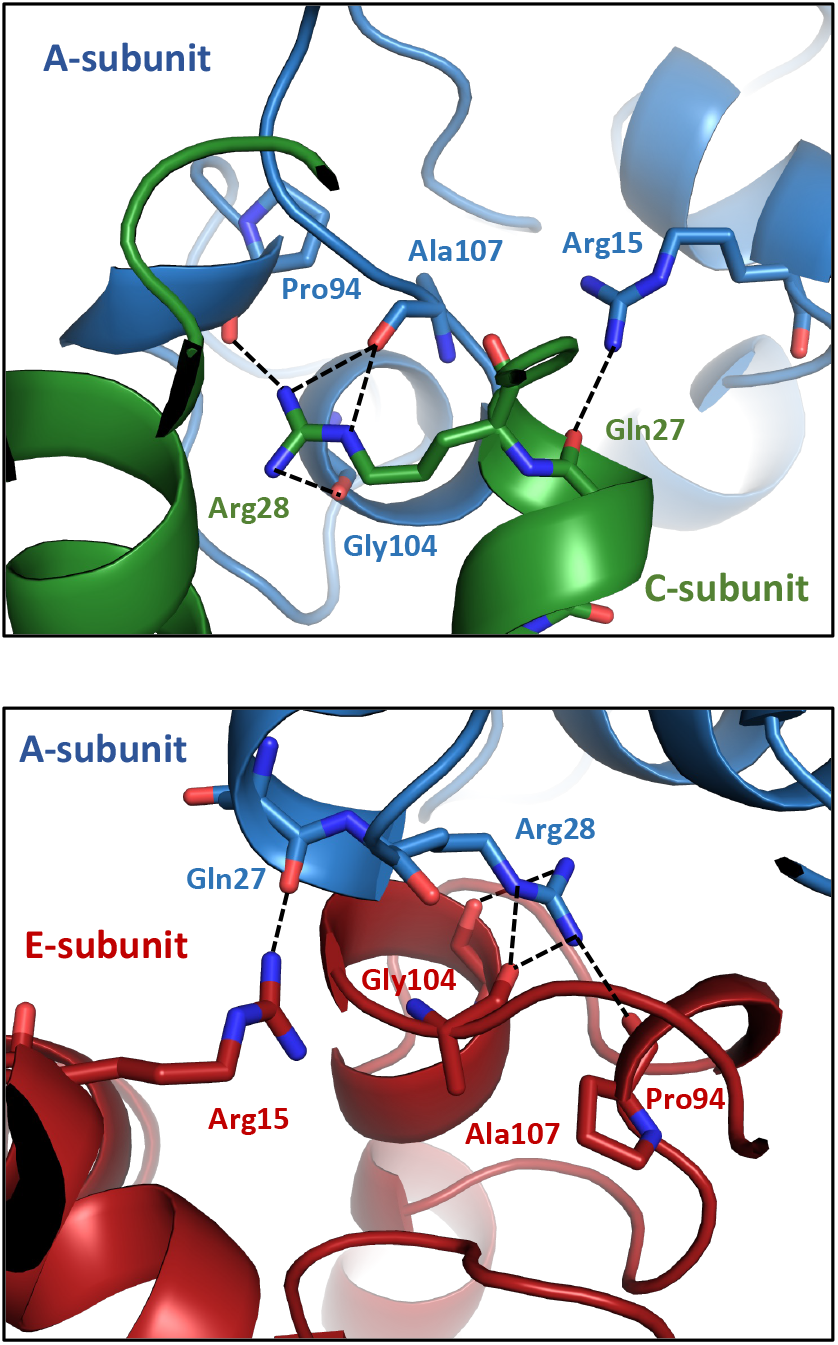
Subunit interface in the hexameric assembly of apo SpNDK. Residues forming H-bonds between the A-C subunits (top), and the A-E subunits (bottom) are shown as sticks.

The crystal structure of the ADP/Vi-bound form of SpNDK was obtained at a resolution of 3.4 Å, with a hexameric assembly present in the asymmetric unit of the crystal. The binding of nucleotide in the presence of vanadate does not induce a major structural change in the monomer (Fig. 4**A**), and both the apo and holo state models could be aligned with a rmsd of 0.32 Å over 120 C_α_.The residues forming the nucleotide binding site are shown in Fig. 4**B** and are either strictly conserved (for Lys9, Arg86, Arg103, Asn118 and His121) or highly conserved (for Phe58, Ile62, Thr92 and Asp124; Fig. 2). One notable exception is Ile116 which is located in the extended *Kpn*-loop of SpNDK. It interacts on one side with the adenine moiety of the bound nucleotide, while Phe58 creates a π-stacking interaction on the other side with the adenine rings. Regarding the vanadate, distances and electron density map suggests that it is bound in a trigonal bipyramidal conformation and is covalently linked to an oxygen atom of the β-phosphate of ADP (Fig. 4**B**), as also observed in the crystal structure of *Borrelia burgdorferi* NDK^18^. However, the limited resolution of our ADP/Vi/NDK complex does not permit to rule out an alternative tetrahedral configuration of vanadate, like the one found in the *S. aureus* enzyme^13^.

**Figure 4.**
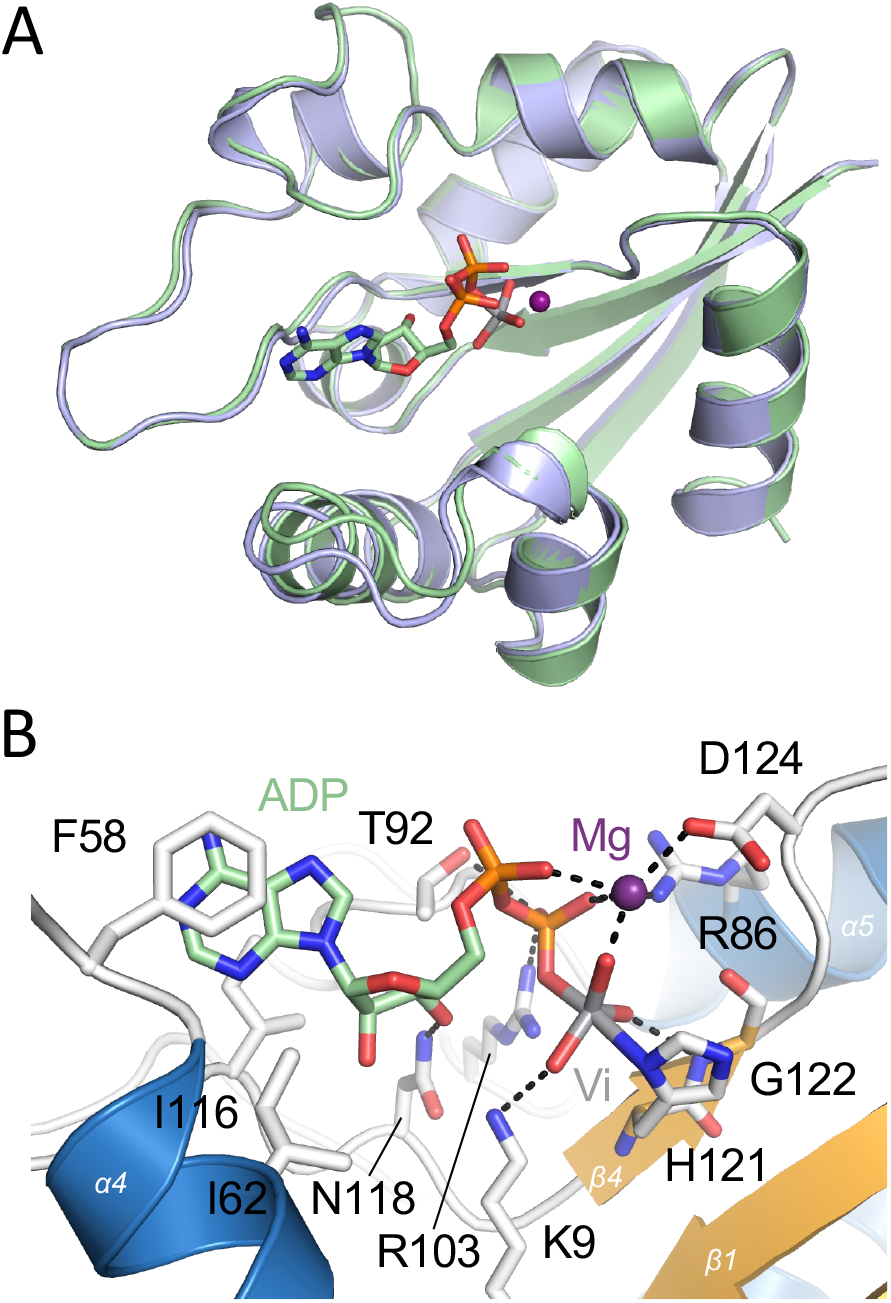
Close-up view of the ADP-Vi binding site of SpNDK. **A**, Structural comparison between apo (blue) and ADP-Vi bound form (green) of SpNDK. **B**, Close-up view of the ADP-Vi binding site of SpNDK. Main chains of interacting residues are in grey sticks. Vanadate (Vi), in trigonal bipyramidal conformation, is covalently bound to the β-phosphate of ADP (green). A magnesium ion (purple) coordinates the ADP phosphates and the vanadate.

The thermal stability of purified SpNDK was assessed by nanoDSF (Fig. S2). The enzyme was remarkably stable with a very high melting temperature (T_m_ ~ 77 °C) which was only moderately further stabilized by nucleotides, Mg-ATP or Mg-GTP, with a T_m_ increased by ~ 2 °C. This is consistent with the moderate impact of nucleotide on the overall structure of SpNDK (Fig. 4**A**). The purified SpNDK was further characterized by size-exclusion chromatography and mass photometry (Fig. 5). Both techniques supported that the main assembly of SpNDK is the hexameric state. However, they revealed unexpectedly that higher oligomeric species were also present in solution, either in concentrated (i.e. size-exclusion, Fig 5**A** and **B**) or diluted (mass photometry, Fig. 5**C**) samples, with the presence of 12-, 18- and presumably 24-Mers of SpNDK. We therefore also solved the cryo-EM structure of hexameric SpNDK at a resolution of 2.47 Å. The map and model are presented in Fig. 6**A** with data collection and refinement statistics in Table S2. As shown in Fig. 6**B**, the model is highly similar to the crystal structure (rmsd of 0.364 over 125 C_α_). Interestingly, if the protein was mostly found in the hexameric state, we could isolate two similar dodecameric assemblies, refined to ~ 4 Å resolution (Fig. 6C and S3). In both assemblies, However, the preferential orientations of the dodecamers in the frozen samples and the limited resolution of the maps (Fig. S4) precluded atomic model refinement. Nonetheless, by fitting the cryo-EM hexamers in both maps, we found that in each assembly, one NDK hexamer interacts with another one at an angle of 70-75°. Although identification of the precise interface between the two hexamers remains challenging, we tentatively proposed several residues–shown as sticks in Fig 6C–that could potentially stabilize the dodecameric form. Additional work will be required to elucidate precisely how the two hexamers, and higher oligomeric states, interact.

**Figure 5.**
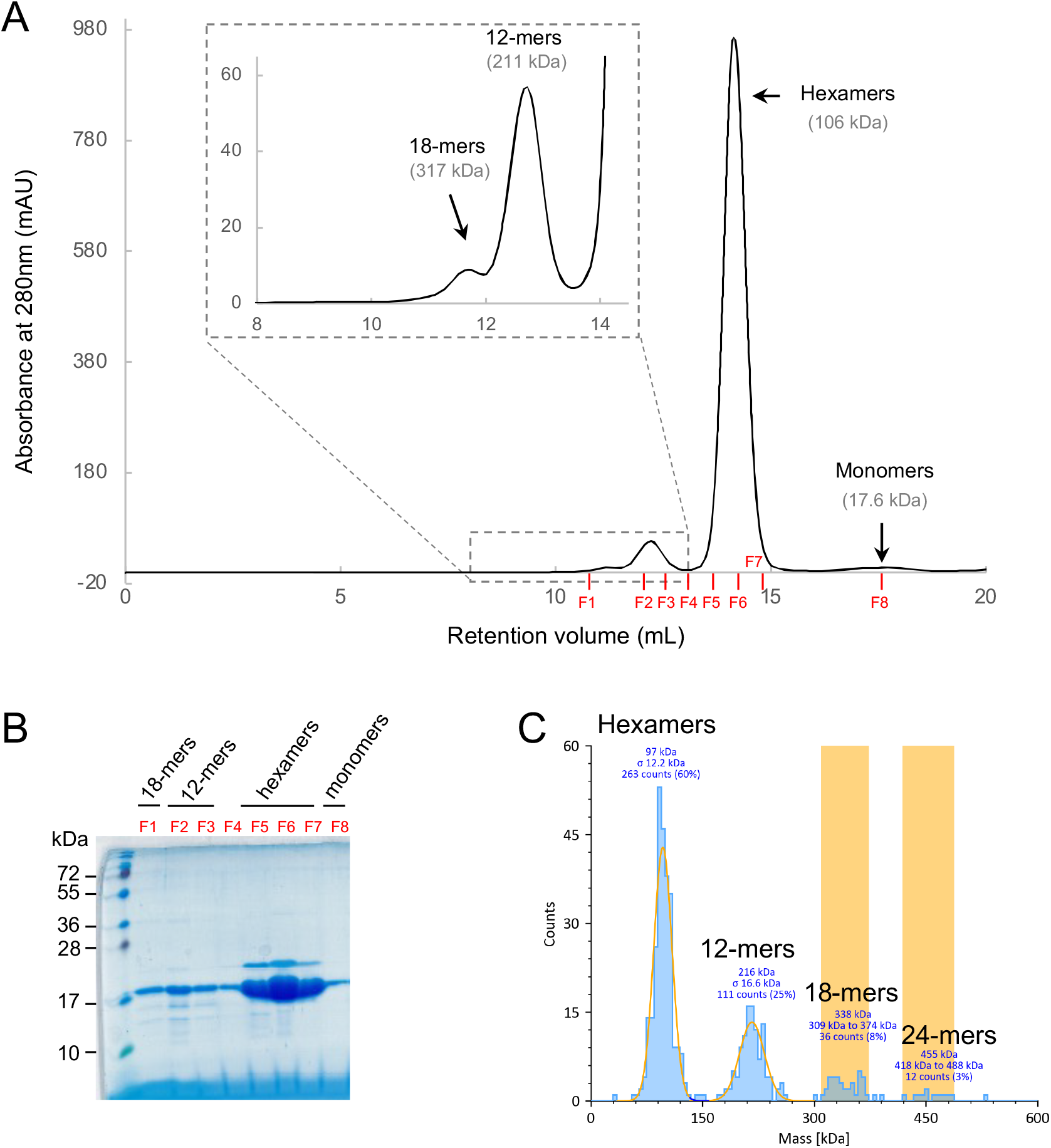
Size-exclusion chromatography and mass photometry of SpNDK. **A**, SEC profile for purified SpNDK. Elution peaks are shown with the estimated molecular weight of each different oligomerization state, deduced from a standard curve. The inset is a magnification of the two peaks corresponding to the 18- and 12-mers. **B**, the collected fractions (F1 to F8) as indicated in red in panel A were resolved on an 18% SDS-PAGE gel. **C**, Mass photometry profile for SpNDK with gaussian peaks (high count numbers) and histograms (low count numbers) shown for the NDK different oligomeric states. The percentage of each oligomeric state is also indicated.

**Figure 6.**
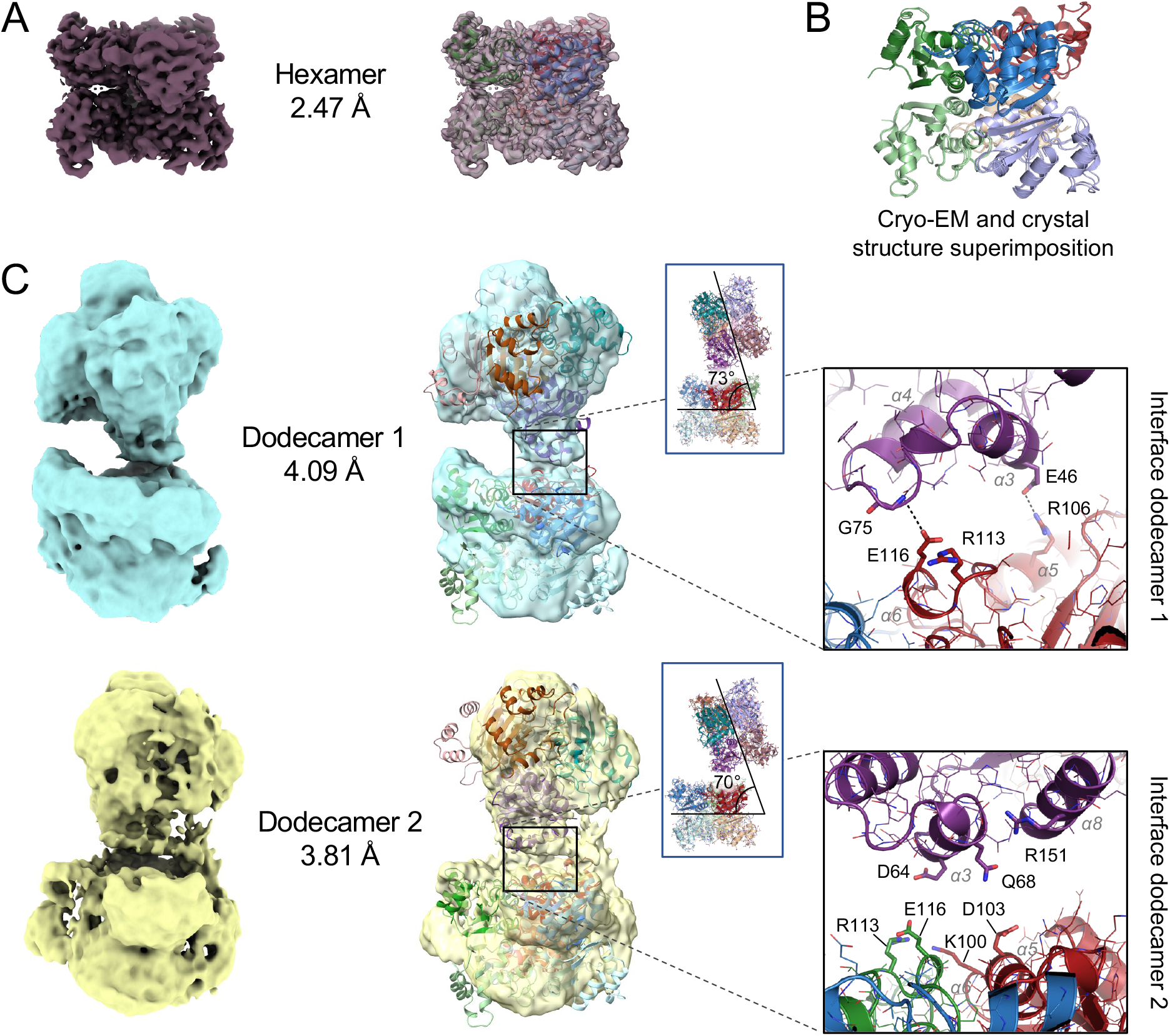
Cryo-EM structures of spNDK. **A**, Cryo-EM map and model fitting of the hexameric NDK. Each monomer is represented in a different colour. **B**, Superimposition of the cryo-EM model (9SLP) with the crystal structure (9SFO) of hexameric NDK, displaying very high similarity (rmsd of 0.364 over 125 C_α_). **C**, Cryo-EM maps and model fitting of the hexameric NDK (9SLP) into dodecamer map 1 (top) and 2 (bottom). Relative position of each NDK hexamer within each dodecamer is shown in insert and a close-up view highlighting residues at the interface between two monomers is displayed on the right. Due to the low resolution of the maps, models could not be refined; residue side chains are therefore as observed in the hexameric form.

## DISCUSSION

In this study, we solved the crystal structure of SpNDK in both the apo state and the ADP-Vi bound state, revealing that the enzyme adopts a canonical hexameric conformation. In many NDKs, this hexameric state is often stabilized by a C-terminal extension that supports the trimer-of-dimers arrangement, as observed in the *B. burgdorferi* enzyme^18^. This extension has also been reported to influence the hexamer stability in human NDKs^19^, and in the *D. discoideum* enzyme when its deletion is combined with a P100S mutation (“Killer of prune” mutation)^20^. The C-terminus of SpNDK is, however, relatively short, a feature shared with some bacterial NDKs that forms exclusively tetramers^11^. Therefore, the hexameric assembly of SpNDK must be stabilized by other regions of the protein. Here, hexamer integrity is notably secured by the *Kpn*-loop, which is highly divergent as compared to other NDK structures, as well as by residues located between β-strands 1 and 2 (Figure S1). In *M. tuberculosis*, the NDK enzyme (MtNDK)^14^ also lacks a C-terminus extension, yet it forms hexamers that are very stable with a melting temperature of ~ 76 °C^14^, similar to what we found for SpNDK. In *M. tuberculosis*, this thermal stability has been attributed to the presence of a salt bridge formed between Arg80 of one subunit and Asp93 of another subunit from a neighboring dimer^21^. Disruption of this salt bridge by the mutation of Asp93 into asparagine prevented the formation of six D93-R80 salt bridges in the whole hexamer, thereby leading to a dramatic T_m_ decrease of ~ 28 °C^21^. Arginine residues are frequently found at subunit-subunit interface^22,23^, and their ability to create up to five H-bonds—in particular with carboxyl groups of the main chain residues—is an efficient strategy used by proteins throughout evolution to increase their stability^24^. In *S. pneumococcus*, the stabilization of the trimer interface within the hexamer involves two arginine residues, Arg15 and Arg28, from one subunit, each forming H-bonds with residues from a neighboring dimer. Arg28 especially contributes to the stabilization of the trimer by forming four H-bonds with two main chain carboxyl groups, including Pro94, from a neighboring subunit. In Drosophila, the mutation of the residue equivalent to Pro94 into Ser (“Killer of prune” mutation) has been shown to cause a substantial reduction in protein thermostability^25^, with the melting temperature decreasing from 71 °C for the wild-type protein to 57 °C for the mutant^26^, while the enzyme activity was unaffected. Despite the large variability in the *Kpn*-loop sequences and in subunit-subunit interactions derived thereof, these results underscore the key role of this loop in the stabilization of the quaternary structure of many hexameric NDKs. SpNDK is a vivid example of this property, although it exhibits some specific features in the residues mediating subunit interactions.

The high thermostability of SpNDK was unexpected as this property is generally reserved to thermophilic bacteria^27,28^. For example, the NDKs of two mesophilic bacteria, *E. coli* and *B. subtilis*, have a melting temperature below 60°C, whereas the NDK of *Vibrio cholerae* exhibits an even lower T_m_ (44°C)^29^. In contrast, the NDK from the thermophilic archaeon *M. thermautotrophicus* has a T_m_ of approximately 80°C^27^. Because NDKs from pathogenic micro-organisms are often secreted and act as virulent factors^7^, they must withstand various harsh environments within host cells^30^. Consequently, it has been proposed that the high stability of some NDKs is an intrinsic property that enables the enzyme to maintain catalytic activity under hostile conditions^11,30,31^. *Streptococcus pneumoniae* can thrive in multiple human organs, and its spreading is notably controlled by macrophages^1^. Although it is currently unknown whether SpNDK can be secreted by *S. pneumoniae*, this bacterium can undergo spontaneously autolysis, releasing its intracellular content into the surrounding milieu^32^. This mechanism is notably responsible for the release of pneumolysin, one of the major virulence factors of *S. pneumoniae*^33^, into infected tissues^34^. Therefore, it is tempting to speculate that the high stability of SpNDK could be advantageous after autolysis, enabling the enzyme to resist host defenses and potentially promoting *S. pneumoniae* infection.

The oligomeric state, in particular for NDKs that assemble into hexamers, is required for the nucleoside diphosphate kinase activity^21^. This is because it stabilizes the *Kpn*-loop at the interface of the trimers-of-dimers assembly^35^, a loop that is directly involved in nucleotide binding^11^. Beyond the canonical hexameric state, a key finding from our study is the ability of SpNDK to form supramolecular structures, suggesting a potential new layer of regulation for the diverse cellular functions of NDKs. Indeed, it is well-known that variation in oligomerization status provides an efficient means to diversify the protein functions within the cell^36^. For example, the B-isoform of human NDK typically assembles as hexamers^37^. However, its ability to bind single strand DNAo has been shown to be linked to the dimeric form of the enzyme^39^. Similarly, purified NDKs from *E. coli* or *V. cholerae* exist in equilibrium between a canonical tetramer and a dimer^29,40^. This equilibrium occurs at physiologically relevant concentrations for *E. coli* enzyme^40^, supporting the possibility of a yet uncharacterized functional role for the dimer. Nucleotide binding shifts this equilibrium *in vitro*: ATP or ADP promotes tetramer formation, indicating that the concentration of nucleotides is a key regulator of the protein oligomeric state^40^.

Finally, some studies have briefly reported the occurrence of NDK dodecamers in plants^41,42^, with one case demonstrating functional activity of this form using blue native gels^41^. Although these observations were not further discussed in these studies, they hint that supramolecular assemblies may also occur in other NDK species.

In conclusion, our work establishes the structural basis of SpNDK oligomerization at atomic resolution and reveals unprecedented supramolecular assemblies. These assemblies could potentially modulate the enzymatic activity or interactions with cellular partners, given the remarkable functional diversity of NDKs and their involvements in various cellular processes. Further studies will determine whether this novel property is conserved across other NDKs and how it may impact their physiological roles.

## Acknowledgements

We acknowledge the European Synchrotron Radiation Facility (ESRF) for providing access to synchrotron radiation facilities, the UAR3033/US001-IECB cryo-EM platform, and the CNRS-IN2P3 computing center (Lyon - France) for providing computing and data-processing resources needed for this work. We thank the contribution of PSF (Protein Science Facility) from SFR Biosciences (CNRS UAR3444). We acknowledge the ISBG (Integrated Structural Biology Grenoble) platform for providing access to the mass photometry system, part of the Grenoble Instruct-ERIC center (ISBG; UAR 3518 CNRS-CEA-UGA-EMBL) within the Grenoble Partnership for Structural Biology (PSB), supported by FRISBI (ANR-10-INBS-0005-02) and GRAL, and financed within the University Grenoble Alpes graduate school (Ecoles Universitaires de Recherche) CBH-EUR-GS (ANR-17-EURE-0003). The financial support of the ANR to J.-M.J. and O.L. (Grant N° 22-CE11-0018) including two PhD contracts to J.K. and P.N. is gratefully acknowledged.

## Materials and Methods

### Plasmid construction

NDK carrying a N-terminal hexa-histidine tag (His-NDK) was amplified by polymerase chain reaction (PCR) from *Streptococcus pneumoniae* genomic DNA using the forward CATATGGAACAAACATTCTTTATCATCAAACCA and reverse primer GGATCCTTAAAACCAAAGAGCAATTTC TCG, containing respectively a NdeI or BamHI restriction site. The NdeI/BamHI digested fragment was ligated into pET28 pre-digested with the same enzymes. The final expression plasmid resulted in pET28-*his*-*ndk* which encodes a 157 amino acid polypeptide including an N-terminal histidine-tag, referred to as His-spNDK in the text.

The same cloning strategy was used to construct spNDK-His carrying a C-terminal hexa-histidine tag, except that the primers for the PCR were 51-CGATACCATGGAACAAACATTCTTTATCATCA AACCA-31, and CGTTACTCGAGAAACCAAAGAGCAATTTCTCG-31, containing respectively a NcoI and XhoI restriction site. *E. coli* BL21(DE3) cells were transformed with the plasmid encoding either His-spNDK or spNDK-His.

### Protein purification

For the SpNDK-his, the NDK protein carrying a C-terminal hexa-histidine tag was produced in *E. coli* BL21(DE3) transformed with pET28-*ndk*-his. Bacteria were grown at 37°C in LB media supplemented with 50 μg.mL^−1^ kanamycin. Protein expression was induced at a DO_600_ of 0.5 by the addition of 1 mM IPTG. After 16h of incubation at 18°C, bacteria were harvested by centrifugation at 6,000 × g at 4°C for 15 min and resuspended in lysis buffer (100 mM HEPES pH 7.4, 150 mM NaCl and 5% (w/v) glycerol). Cells were lysed by sonication with 10 cycles of 30 s with 30 s interval between each pulse and the lysate was centrifuged at 10,000 × g for 30 min at 4°C. The supernatant containing spNDK-his was loaded onto a Nickel affinity column (HisTrap™ Fast Flow, Cytiva) equilibrated with lysis buffer containing 40 mM imidazole. Bound proteins were eluted in lysis buffer supplemented with 400 mM imidazole. The eluate containing spNDK-his was loaded onto a size-exclusion chromatography column (Superdex™ 200 Increase 10/300 GL, Cytiva) equilibrated with 50 mM HEPES pH 7.4, 50 mM NaCl and 1 mM EDTA. Fractions of interest were pooled and concentrated to 10 mg.mL^−^1 using an Amicon® Ultra-4 centrifugal device (Millipore®) with a molecular weight cut-off of 3 kDa.

For his-SpNDK, the NDK protein carrying a N-terminal hexahistidine tag was produced in *E. coli* BL21(DE3) transformed with pET28-*his-ndk*. Bacteria were grown at 37°C in 500 mL of LB media supplemented with 50 μg/mL kanamycin for 4-6h until a DO_600_ of 0.6 was reached. The culture was then induced with 0.7 mM IPTG for 1h at 25°C, and centrifuged at 7000 *g* for 10 min at 4°C. The bacteria were washed with 35 mL of buffer 50 mM Tris pH 8.0 and 5 mM MgCl_2_, and finally resuspended in 50 mM Tris pH 8.0, 5 mM MgCl_2_, 1 mM DTT, CLAPA protease inhibitors and 1 mM DNase. The cells were lysed by sonication with 6 cycles of 20 sec and 20 sec of interval between each impulsion. The lysate was then centrifuged at 150,000 g for 1h at 4 °C and 2 mL of Ni-NTA resin previously equilibrated with 50 mM HEPES pH 7.4, 100 mM NaCl and 10 mM imidazole were added to the supernatant. The resin was then washed 5 times with 3 mL of the same buffer, and bound proteins were eluted with twice 2 mL of buffer supplemented with 300 mM imidazole. The proteins were centrifuged at 15,000 g for 15 min at 4°C to remove the aggregates before injection onto a Superdex 200 Increase 10/300 GL column, previously equilibrated with 50 mM HEPES pH 7.4 and 100 mM NaCl.

### Protein crystallization and structure determination of the NDK structures

His-SpNDK was crystallized at a concentration of 17.5 mg/mL by mixing 0.5 µL with an equal volume of precipitation reagent in an MRC 2-drop plate (Molecular Dimensions), using the liquid-handling robot Mosquito (TTP). The reservoir volume was 70 µL. Crystals of the apo protein were obtained at 20°C in 0.2 M calcium chloride dihydrate, 0.1 M HEPES sodium pH 7.5 and 28% (v/v) PEG 400. Crystals were directly flash-frozen in liquid nitrogen and diffracted remotely on beamline ID30A-3 (MASSIF-3) at the ESRF synchrotron. For the ADP-bound structure, SpNDK-His was used. Crystals were obtained by mixing an equal volume of protein at 18 mg/mL and reservoir solution composed of 24% (v/v) PEG 3350, 100 mM MES pH 7.0 in a final volume of 1.7 µL. Crystallization was performed at 20°C by the hanging drop vapour diffusion method in 24-well XRL plates (Molecular Dimensions), over a reservoir solution of 500 µL. Ligands were soaked into the crystal-containing drop by adding a solution of orthovanadate (5 mM final) previously heated at 90°C for 5 min and MgADP (5 mM final). After 5 min, crystals were cryoprotected with 15% of ethylene glycol before being flash frozen in liquid nitrogen and remote diffraction on beamline ID23-2 (ESRF).

For both datasets, images were indexed and integrated with iMosflm, then scaled and merged with Aimless^43^. The apo structure was solved by molecular replacement using Phaser^44^ with the NDK model from *Staphylococcus aureus* (PDB 3Q8Y) as the search probe, after removal of ligands. The apo model was then used to solve the liganded-NDK structure. The unit cell contains one monomer per asymmetric unit for the apo structure and six for the ADP-vanadate structure. The models were refined with cycles of restrained refinement with REFMAC5^45^ and manual building in Coot^46^. The final structures were validated using Procheck^47^ and Rampage^48^. Data collection and refinement statistics are listed in Table S1.

### Nano Differential Scanning Fluorimetry

The NanoDSF experiment was performed using a Prometheus NT.48 instrument (NanoTemper Technologies) in a buffer composed of 50 mM HEPES pH 7.4, and 50 mM NaCl. The C-terminal His-tag of his-SpNDK was removed using the Thrombin CleanCleave Kit (Sigma) and the tag-free protein was used at 32 µM (0.5 mg/mL). When specified, ATP or GTP was added at a final concentration of 4 mM in the presence of 5 mM MgCl_2_. Proteins were loaded into NanoDSF grade standard capillaries (NanoTemper Technologies), and exposed to a thermal gradient ranging from 20°C to 95°C with a temperature increase of 1°C every minute. Samples were excited at 280 nm, and the emission of the tryptophan and tyrosine fluorescence was measured at 330 nm and 350 nm using a dual-UV detector. Data were analysed using PR stability analysis software (NanoTemper Technologies).

### Mass determination by mass photometry

All measurements were acquired using a commercial Refeyn OneMP mass photometer. The buffer solution (50 mM HEPES pH 7.4, and 100 mM NaCl) was filtered onto an Amicon Ultra-4 spin centrifugal filter device with a molecular weight cut-off of 30 kDa (Millipore®). The standard native marker (NativeMark™ LC0725, ThermoFisher Scientific) was used for calibration applying three reference points with a maximum calibration error of 5%. The his-SpNDK protein was diluted at 200 nM in filtered buffer. For analysis, 15 μL of filtered buffer was added to an empty well on the cover slide, along with 5 µL of the pre-diluted protein solution. Measurements were acquired using the Acquire MP v2.5.0 programme and ratiometric images (data) were analysed using the Refeyn Discover MP software (Refeyn Ltd.).

### Cryo-EM sample preparation and data collection

A volume of 3 µL of purified his-NDK at 0.25 mg/ml, was applied onto a Quantifoil Au-200-mesh R1.2/1.3 grid previously glow discharged with ELMO system (Cordouan Technologies) for 40 s with 2.7 mA current. Excess solution was blotted for 4 s and vitrification was performed in liquid ethane using a FEI Vitrobot Mark IV (Thermo Fischer Scientific) at 4°C and 100% humidity.

The grid was imaged using a Glacios TEM (Thermo Fischer Scientific) equipped with a Falcon 4i direct electron detector and a Selectris X energy filter. The sample was imaged at a nominal magnification of 165,000x corresponding to a calibrated pixel size of 0.694 Å. A total of 3,184 movie frames with an electron dose of 59.5 e^−^/Å^2^ over 42 frames were collected using EPU software (Thermofisher Scientific) with a defocus range of −0.5 to −1.7 µm (steps of −0.2 µm).

### Cryo-EM image processing

Data analysis was carried out using cryoSPARC v4.6.0^49^. Movies were corrected for gain reference, motion-corrected using Patch motion correction and the contrast transfer function (CTF) was determined using Patch CTF. For NDK hexamer, a total of 3,231,210 particles were picked using blob picker and subsequently extracted. After two rounds of 2D classification to remove 2D class averages with less resolved features or presenting particles in close contact, the selected 716,048 particles were used to build five *ab initio* maps. The best isotropic map was used as the initial reference to calculate a new map from the 716,048-particle dataset using hetero-refinement classification. The hexamer map was further improved using non-uniform refinement by imposing D3 symmetry yielding a map with resolution at 2.47 Å. For NDK dodecamer, a total of 1,653,163 particles were picked using automated blob picker with an elliptical shaped blob. After two rounds of 2D classification to remove 2D class averages with less resolved features or presenting a single hexamer, the selected 148,143 particles were used to generate three *ab initio* maps. Two maps were obtained with a dodecamer organisation. The corresponding particles were separated in five 3D classes using 3D classification leading to two alternate conformations. The first and second maps (dodecamer 1 and 2), comprising 62,236 and 51,255 particles, respectively, were refined independently using non-uniform refinement without imposing symmetry. The final resolutions of dodecamer 1 and 2 were 4.09 Å and 3.81 Å, respectively.

## Cryo-EM model building

A first monomer model was generated using the ‘Predict and Build’ tool implemented in Phenix^50^ from the cryo-EM hexamer data. This model was refined using Isolde^51^ and Phenix. Structure validation was performed with MolProbity program^52,53^. Data collection and refinement statistics are listed in Table S2.

## Data availability

The coordinates and structure factors for the NDK crystal models have been deposited in the Protein Data Bank (PDB) under accession numbers 9RVW (apo form) and 9SFO (ADP-Vi complex). The cryo-EM atomic model and maps have been deposited in the PDB and Electron Microscopy Data Bank (EMDB) under the accession codes 9SLP and EMD-55011 for the NDK hexamer, and EMD-55334 and EMD-55333 for the NDK dodecamer 1 and 2 maps, respectively.

## Figure legends

**Figure S1. Secondary structures and intermolecular contacts for selected NDKs**. The secondary structures are shown above the sequences and were derived from the 3D structures of SpNDK (PDB 9RVW), *Mycobacterium tuberculosis* (1K44), *Borrelia burgdorferi* (4DI6), *Pyrobaculum aerophilum* (1XQI) and *Escherichia coli* (2HUR). The four first NDKs assemble as hexamers (top left panel) and the last NDK, from *E. coli*, forms tetramers (top right panel), with the subunit arrangement indicated. Residues adopting alternative conformations are highlighted by a grey star above the sequence. The accessibility (acc), hydropathy scales (hyd), and intermolecular contacts (bottom lines underneath the sequences) are shown. The intermolecular contacts between the A subunit (query) and the other subunits from the biological assembly (e.g. subunits C, B and E for SpNDK) are indicated. For intermolecular contacts < 3.2 Å, the letter is in red, and for contacts between 3.2-5.0 Å, the letter is in black. For SpNDK and *M. tuberculosis* NDK, the residues involved in the main interactions are indicated by double-sided arrows (e.g. between R_15_ in the A-subunit of SpNDK and Q_27_ in the C-subunit) with interactions between A- and C-subunits, and A- and E-subunits, in green and firebrick colours, respectively. Green numbers 1 below the *B. burgdorferi* sequence indicate the formation of a disulphide bond. The figure was drawn with ENDscript 2.0^54^.

**Figure S2. Thermal stability of NDK measured by NanoDSF**. SpNDK was preincubated 20 min at 22°C with MgATP (red curve), MgGTP (green curve) or without any ligand (blue curve) prior to gradually increasing the temperature at 1°C.min^−1^. The fluorescence ratio (F_350nm_/F_330nm_) and its first derivative are shown in the top and bottom panels, respectively. The inflection point of the derivatives, corresponding to the melting temperatures (T_m_), are reported in the inset of the bottom panel.

**Figure S3. Cryo-EM processing workflow for NDK hexamer and dodecamers**. After patch motion correction and CTF estimation, hexameric NDK particles were picked with a circular blob picker. Following rounds of 2D classification, five *ab initio* models were generated, with the best map serving as reference for heterogenous refinement. The resulting hexamer map was further refined with D3 symmetry, yielding a final map at 2.47 Å resolution (FSC_0.143_). For dodecameric NDK, particles were picked with an elliptical blob picker. After two rounds of 2D classification, three *ab initio* maps were generated. A subsequent 3D classification produced two conformations (dodecamer 1 and 2), which were refined separately using non-uniform refinement to 4.09 Å and 3.81 Å resolution (FSC_0.143_).

**Figure S4. Cryo-EM analysis of NDK**. Viewing direction distribution plots of particles for NDK hexamer (**A**), dodecamer 1 (**C**) and dodecamer 2 (**E**). Most frequent views are colored in red. Final cryo-EM map of NDK hexamer (**B**), dodecamer 1 (**D**) and dodecamer 2 (**F**) colored according to the local resolution.

Values in parentheses indicate the outer resolution bin. Reflection data and refinement statistics are as reported by Aimless^43^ and REFMAC5^45^, respectively. Ramachandran statistics are from Rampage^48^.

**Table S1.**
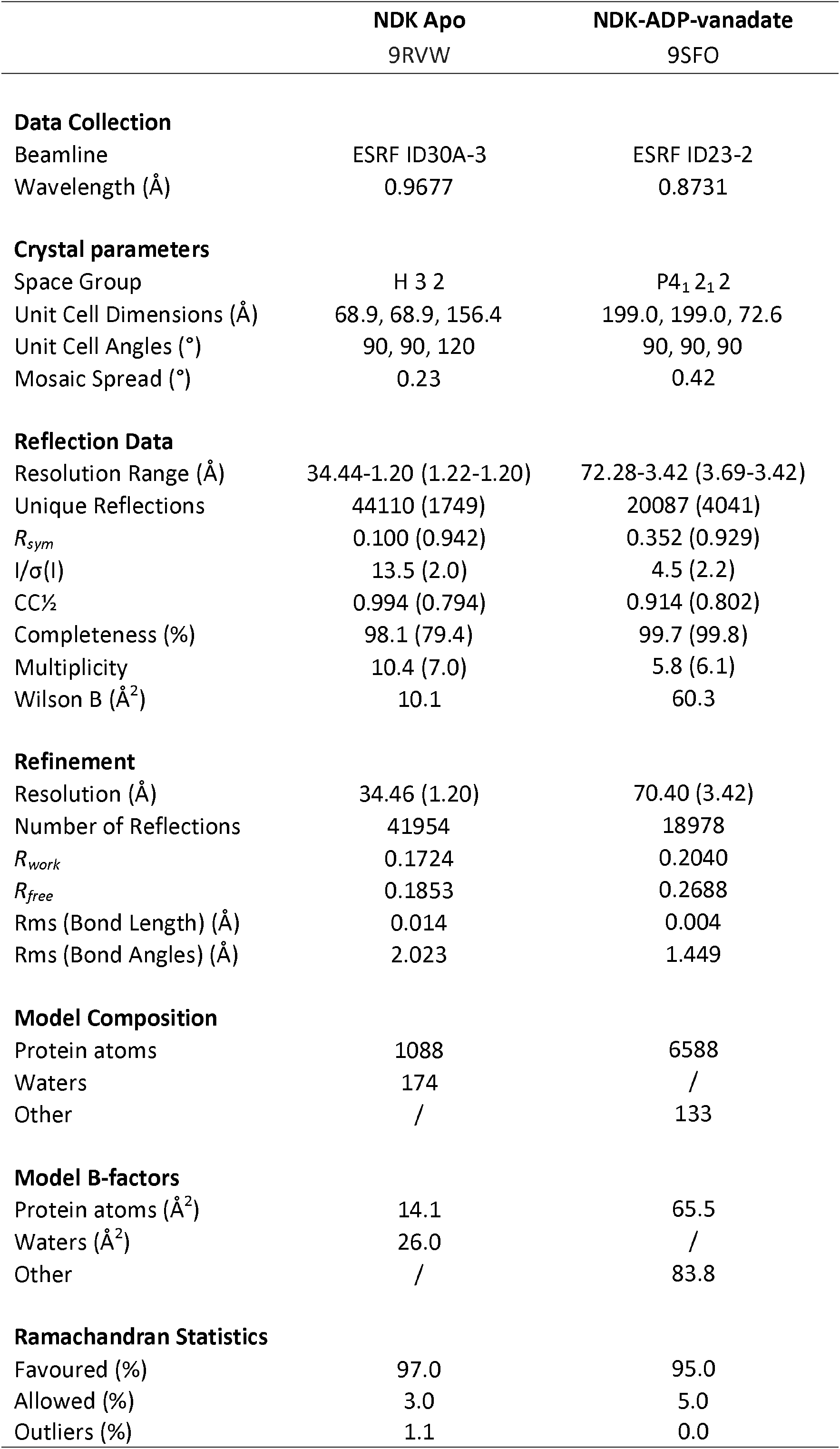
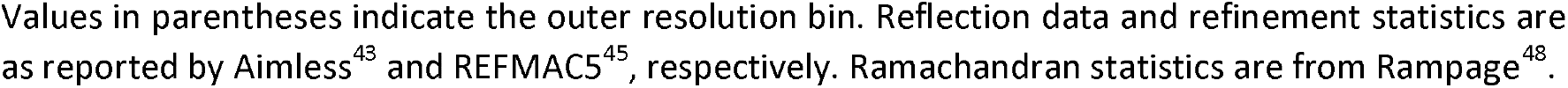
X-ray data and refinement statistics of SpNDK.

|  | **NDK Apo**<br>9RVW | **NDK-ADP-vanadate**<br>9SFO |
| --- | --- | --- |
| **Data Collection** |  |  |
| Beamline | ESRF ID30A-3 | ESRF ID23-2 |
| Wavelength (Å) | 0.9677 | 0.8731 |
| **Crystal parameters** |  |  |
| Space Group | H 3 2 | P4<sub>1</sub> 2<sub>1</sub> 2 |
| Unit Cell Dimensions (Å) | 68.9, 68.9, 156.4 | 199.0, 199.0, 72.6 |
| Unit Cell Angles (°) | 90, 90, 120 | 90, 90, 90 |
| Mosaic Spread (°) | 0.23 | 0.42 |
| **Reflection Data** |  |  |
| Resolution Range (Å) | 34.44-1.20 (1.22-1.20) | 72.28-3.42 (3.69-3.42) |
| Unique Reflections | 44110 (1749) | 20087 (4041) |
| $R_{sym}$ | 0.100 (0.942) | 0.352 (0.929) |
| $I/\sigma(I)$ | 13.5 (2.0) | 4.5 (2.2) |
| CC<sub>1/2</sub> | 0.994 (0.794) | 0.914 (0.802) |
| Completeness (%) | 98.1 (79.4) | 99.7 (99.8) |
| Multiplicity | 10.4 (7.0) | 5.8 (6.1) |
| Wilson B (Å<sup>2</sup>) | 10.1 | 60.3 |
| **Refinement** |  |  |
| Resolution (Å) | 34.46 (1.20) | 70.40 (3.42) |
| Number of Reflections | 41954 | 18978 |
| $R_{work}$ | 0.1724 | 0.2040 |
| $R_{free}$ | 0.1853 | 0.2688 |
| Rms (Bond Length) (Å) | 0.014 | 0.004 |
| Rms (Bond Angles) (Å) | 2.023 | 1.449 |
| **Model Composition** |  |  |
| Protein atoms | 1088 | 6588 |
| Waters | 174 | / |
| Other | / | 133 |
| **Model B-factors** |  |  |
| Protein atoms (Å<sup>2</sup>) | 14.1 | 65.5 |
| Waters (Å<sup>2</sup>) | 26.0 | / |
| Other | / | 83.8 |
| **Ramachandran Statistics** |  |  |
| Favoured (%) | 97.0 | 95.0 |
| Allowed (%) | 3.0 | 5.0 |
| Outliers (%) | 1.1 | 0.0 |

Values in parentheses indicate the outer resolution bin. Reflection data and refinement statistics are as reported by Aimless<sup>43</sup> and REFMAC5<sup>45</sup>, respectively. Ramachandran statistics are from Rampage<sup>48</sup>.

**Table S2.** Cryo-EM data collection and refinement statistics for spNDK.

|  | **NDK hexamer**<br>PDB 9SLP / EMD-55011 | **NDK dodecamer 1**<br>EMD-55334 | **NDK dodecamer 2**<br>EMD-55333 |
| --- | --- | --- | --- |
| **Data Collection and Processing** |  |  |  |
| Microscope | TFS Glacios | TFS Glacios | TFS Glacios |
| Camera | TFS Falcon 4i | TFS Falcon 4i | TFS Falcon 4i |
| Energy Filter | Selectris X | Selectris X | Selectris X |
| Magnification | 165k | 165k | 165k |
| Voltage (kV) | 200 | 200 | 200 |
| Electron exposure (e<sup>-</sup>/Å<sup>2</sup>) | 59.5 | 59.5 | 59.5 |
| Defocus range (μm) | -0.5 to -1.7 | -0.5 to -1.7 | -0.5 to -1.7 |
| Pixel size (Å) | 0.694 | 0.694 | 0.694 |
| Symmetry imposed | D3 | C1 | C1 |
| Initial particle images (no.) | 2,612,955 | 1,653,163 | 1,653,163 |
| Final particle images (no.) | 531.268 | 62.236 | 51.255 |
| Map resolution (Å) | 2.47 | 4.09 | 3.81 |
| FSC threshold | 0.143 | 0.143 | 0.143 |
| Map resolution range (Å) | 2.138 - 33.080 | 1.448 - 64.756 | 1.448 - 59.343 |
| **Refinement** |  |  |  |
| Model resolution (Å) | 2.59 |  |  |
| FSC threshold | 0.5 |  |  |
| Map sharpening B factor (Å<sup>2</sup>) | 86.97 |  |  |
| **Model Composition** |  |  |  |
| Non-hydrogen atoms | 6522 |  |  |
| Protein residues | 822 |  |  |
| Other | 0 |  |  |
| **R.m.s. deviations** |  |  |  |
| Bond lengths (Å) | 0.002 |  |  |
| Bond angles (°) | 0.442 |  |  |
| **Validation** |  |  |  |
| MolProbity score | 1.24 |  |  |
| Clashscore | 4.07 |  |  |
| Poor rotamers (%) | 0 |  |  |
| **Ramachandran Statistics** |  |  |  |
| Favoured (%) | 97.8 |  |  |
| Allowed (%) | 2.2 |  |  |
| Outliers (%) | 0 |  |  |

## Notes

### Competing Interest Statement

The authors have declared no competing interest.

