## Supplementary figures and images for "Quaternary structures of *Streptococcus pneumoniae* Nucleoside Diphosphate Kinase: from hexamers to supramolecular assemblies"

### Fig S1-S4

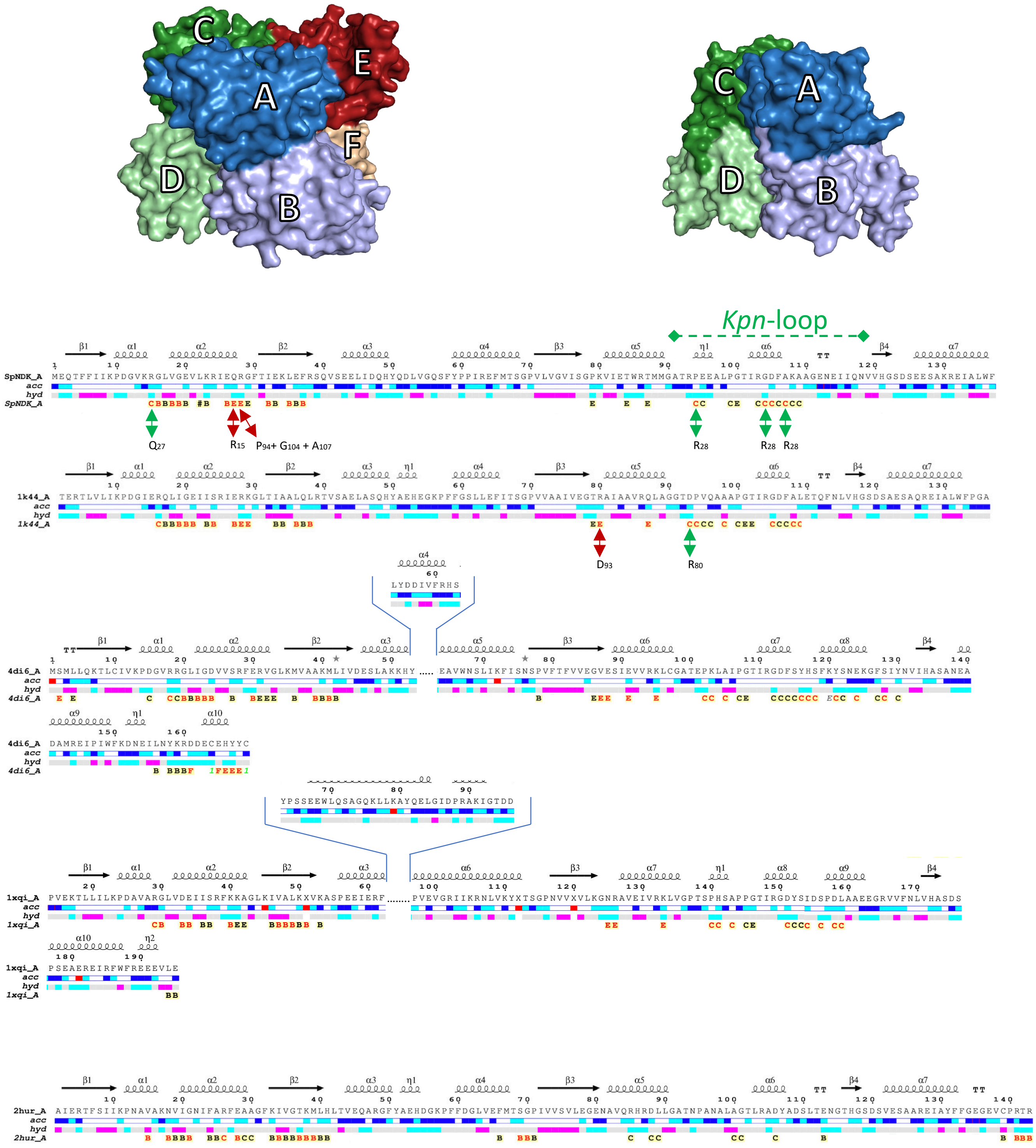

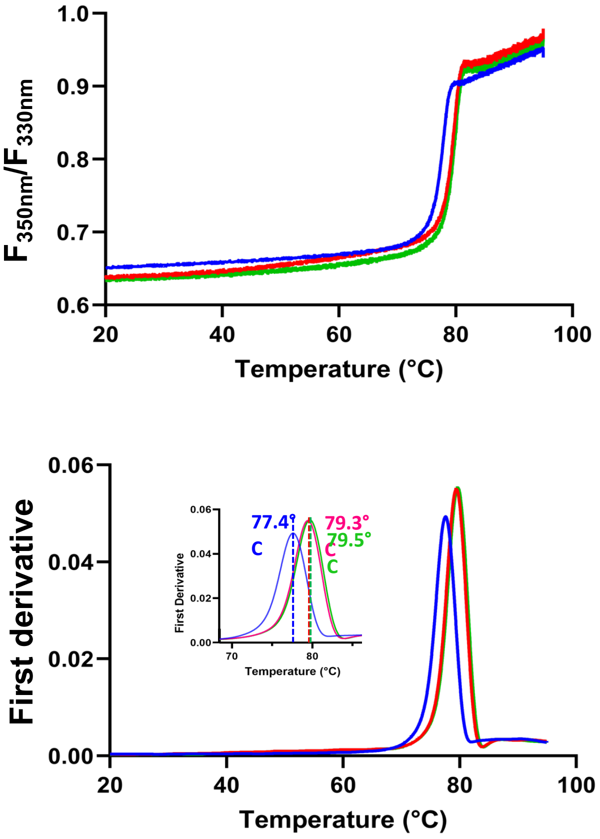

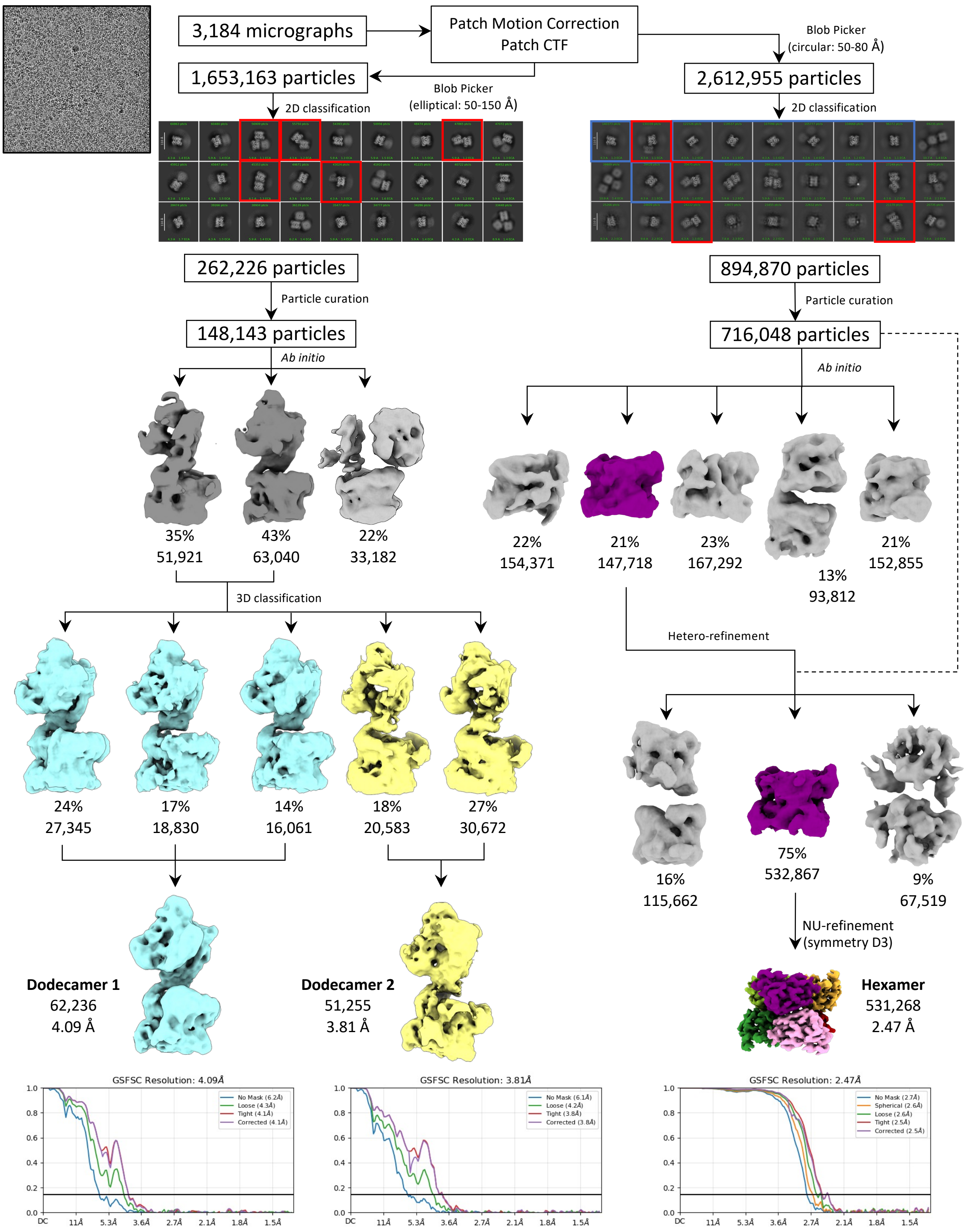

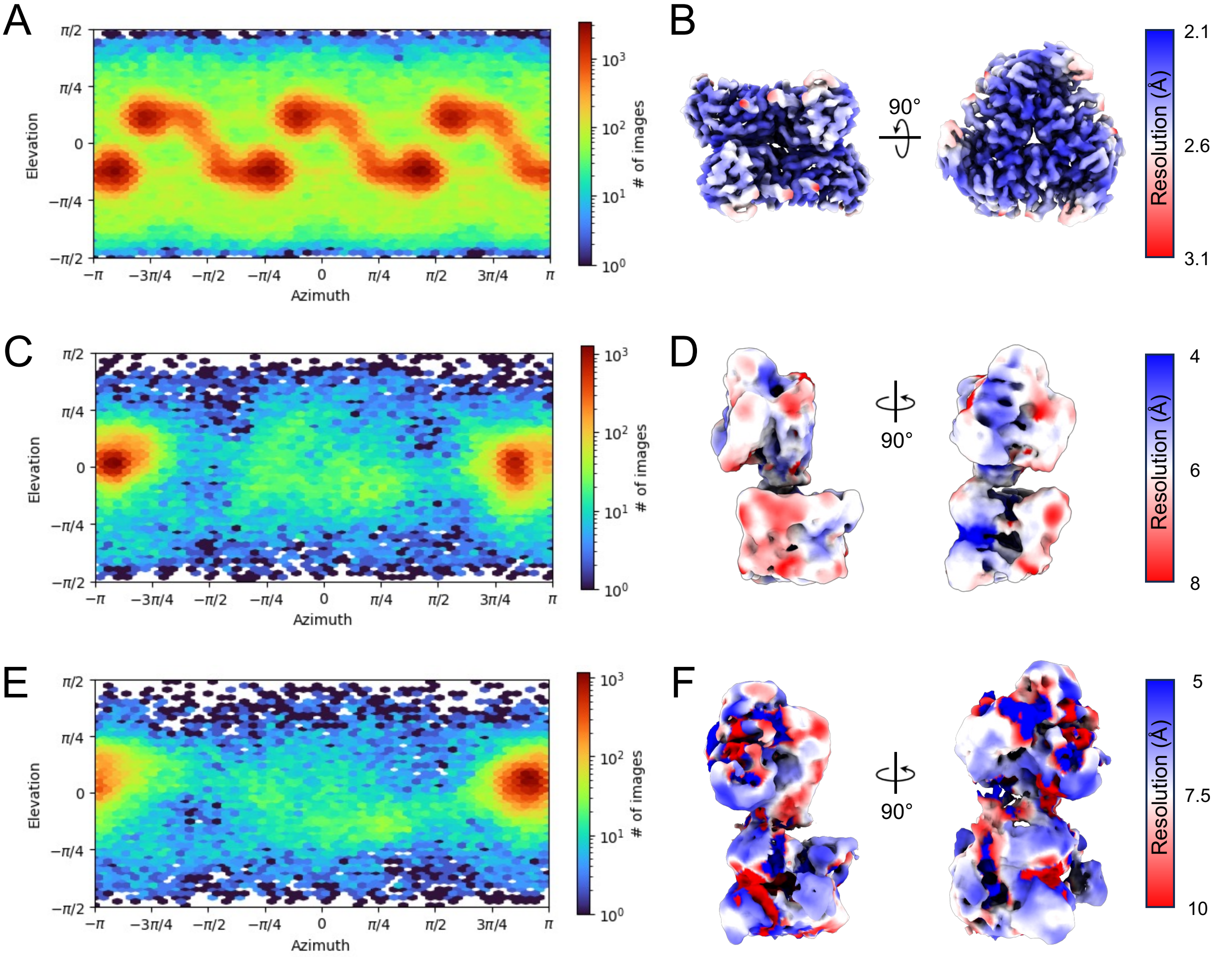
